# *Pituitary Tumor Transforming Gene 1* orchestrates gene regulatory variation in mouse ventral midbrain during aging

**DOI:** 10.1101/2020.05.14.096156

**Authors:** Yujuan Gui, Mélanie H. Thomas, Pierre Garcia, Mona Karout, Rashi Halder, Alessandro Michelucci, Heike Kollmus, Cuiqi Zhou, Shlomo Melmed, Klaus Schughart, Rudi Balling, Michel Mittelbronn, Joseph H. Nadeau, Robert W. Williams, Thomas Sauter, Manuel Buttini, Lasse Sinkkonen

## Abstract

**Background:** Dopaminergic neurons in the midbrain are of particular interest due to their role in diseases such as Parkinson’s disease and schizophrenia. Genetic variation between individuals can affect the integrity and function of dopaminergic neurons but the DNA variants and molecular cascades modulating dopaminergic neurons and other cells types of ventral midbrain remain poorly defined. Three genetically diverse inbred mouse strains — C57BL/6J, A/J, and DBA/2J — differ significantly in their genomes (~7 million variants), motor and cognitive behavior, and susceptibility to neurotoxins.

**Results:** To further dissect the underlying molecular networks responsible for these variable phenotypes, we generated RNA-seq and ChIP-seq data from ventral midbrains of the 3 mouse strains. We defined 1000–1200 transcripts that are differentially expressed among them. These widespread differences may be due to altered activity or expression of upstream transcription factors. Interestingly, transcription factors were significantly underrepresented among the differentially expressed genes, and only one TF, *Pttg1*, showed significant differences among all strains. The changes in *Pttg1* expression were accompanied by consistent alterations in histone H3 lysine 4 trimethylation at *Pttg1* transcription start site. The ventral midbrain transcriptome of three-month-old C57BL/6J congenic *Pttg1^-/-^* mutants was only modestly altered, but shifted towards that of A/J and DBA/2J in nine-month-old mice. Principle component analysis identified the genes underlying the transcriptome shift and deconvolution of these bulk RNA-seq changes using midbrain single cell RNA-seq data suggested that the changes were occurring in several different cell types, including neurons, oligodendrocytes, and astrocytes.

**Conclusion:** Taken together, our results show that *Pttg1* contributes to gene regulatory variation between mouse strains and influences mouse midbrain transcriptome during aging.

## Background

Two populations of dopaminergic neurons (DAns) in ventral midbrain are of translational interest. One group resides in substantia nigra (SN) controlling motor function, while the other is in ventral tegmental area (VTA) and associated with cognitive function (1). Many human phenotypes, such as differences in motor learning (2) or in disease susceptibility to schizophrenia and Parkinson’s disease (PD), are linked to DAns and modulated by genetic variation regulating dopaminergic circuits (3, 4). Interestingly, recent work has established that most genetic variants associated with human traits and diseases are localized in noncoding genome and significantly enriched in cell type-specific gene regulatory regions (5). Indeed, it has been suggested that most complex traits are explained by cumulative effects of numerous *cis-* and *trans*-regulatory variants that individually contributes to relatively small phenotypic effects (6). In particular, peripheral master regulators such as transcription factors (TFs) with tens to hundreds of target genes could be mediating a lot of gene regulatory variation through *trans*-effects while their own expression is altered by local *cis*-variants.

Mouse and human brains share large similarities in dopaminergic circuits and related gene expression, making mouse an ideal model system for neuroscience (1, 7). Three mouse strains, C57BL/6J, A/J, and DBA/2J, are frequently used biology and show phenotypic differences in their dopaminergic circuits. For example, C57BL/6J has the highest motor activity and sensitivity to addiction (8–12), and its dopamine levels in ventral midbrain are increased compared to the other strains (13). Moreover, the strains respond differently to PD toxins such as methyl-4-phenyl-1,2,3,6-tetrahydropyridine (MPTP), drawing parallels with varied susceptibility to PD in human population (14). Mouse models are also a fundamental step to study genetic aspects of the brain, with 90% of mouse genes being identical to human genes (15). Similar to a typical human genome that differs from the reference genome by approximately 5 million variants (16), these mouse strains are collectively segregated by around 7 million variants. These characteristics make the mouse an interesting model to study genetic factors and extent of gene regulatory variation in connection to ventral midbrain and dopaminergic circuits.

Here we aimed to elucidate gene regulatory variation underlying the known phenotypic differences within mouse midbrains (8–12) by using a comparative functional genomics approach focusing on transcriptomic and epigenomic analysis of C57BL/6J, A/J, and DBA/2J strains. We identify significant differences between midbrains of the mouse strains with over 1000 genes showing altered expression levels in each comparison. To delineate whether these changes are due to regulatory variation associated with TFs, we looked at which TFs have altered expression. Suprisingly, TFs are significantly under-represented among the altered genes with only *Pttg1 (Pituitary Tumor Transforming Gene 1)* showing significant changes between all three strains. Deletion of *Pttg1* alone is not sufficient to cause major midbrain gene expression changes in young mice, but does lead to substantial transcriptomic shift during aging, resembling the differences distinguishing C57BL/6J from A/J and DBA/2J strains. The changes induced by loss of *Pttg1* are not limited to any specific cell type but instead appear to affect multiple different cell types of the ventral midbrain. Our findings implicate *Pttg1* in the transcriptomic control of the midbrain during aging, and suggest it could contribute to the gene regulatory variation, and possibly also phenotypic variation, between mouse strains.

## Results

### Midbrain transcriptomes are significantly different between common mouse strains

To investigate genetic background driving gene expression differences in ventral midbrain, we performed transcriptomic and epigenomic analyses on isolated ventral midbrains containing SN and VTA from three genetically diverse mouse strains, C57BL/6J, A/J, and DBA/2J (Figure 1A). For transcriptomic profiling, midbrains from 36 individual 3-month old mice were analysed by RNA-seq, corresponding to 12 mice (6 males and 6 females) from each strain. For epigenomic analysis, the enrichment of histone H3 lysine 4 trimethylation (H3K4me3), an established marker of open transcription start sites (TSS) (17–19), was analysed by ChIP-seq from dissected ventral midbrain of 6 individual 3-month old mice (2 males from each strain).

**Figure 1.**
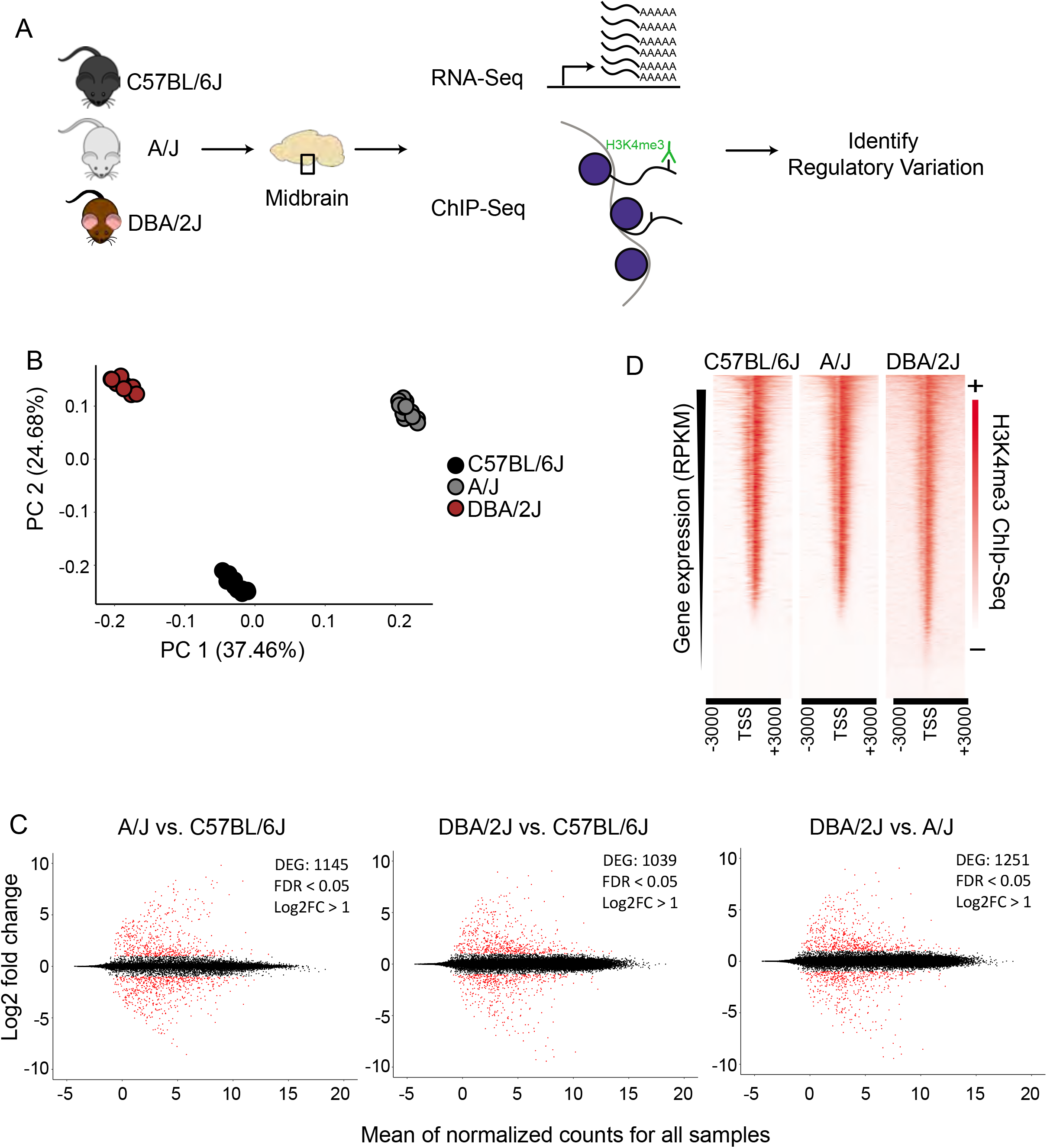
Functional genomics profiling of isolated midbrains of 3-month-old C57BL/6J, A/J and DBA/2J mice. A. Schematic representation of the experimental set-up. The ventral midbrains of C57BL/6J, A/J and DBA/2J, dissected using anatomical landmarks directly after mouse euthanasia, were used for RNA-seq and ChIP-seq. B. Principle component analysis showing transcriptome level differences in the midbrains of the three strains. The individual mice are indicated with black (C57BL/6J), grey (A/J), or brown (DBA/2J) circles. No gender bias was observed. C. Pairwise comparisons showing DEGs in the midbrains of the three strains. MA plots from left to right: A/J vs. C57BL/6J, DBA/2J vs. C57BL/6J, and DBA/2J vs. A/J. The analysis was done by DEseq2 using ashr shrinkage. The x-axis represents the mean of normalized counts for all replicates and the y-axis represents the log_2_-fold change. Each dot represents one gene. Genes with FDR<0.05 and log_2_-fold change (log_2_FC)>1 are indicated in red and referred to as DEGs. D. H3K4me3 ChIP-seq signal with corresponding gene expression levels as measured by RNA-seq. The intensity of H3K4me3 ChIP-seq signals are plotted in a window of 3 kb upstream and downstream of the TSS and within-sample normalization was applied. The genes are ranked based on gene expression levels (RPKM) from highest to lowest.

A principle component analysis (PCA) of RNA-seq data could clearly separate the samples according to strain of origin (Figure 1B), suggesting significant differences exist at the transcriptomic level between ventral midbrains. Indeed, a pair-wise comparison of the individual of strains to each other revealed a significant (FDR<0.05) change in expression with a log_2_-fold change (log_2_FC) higher than 1 for more than 1000 genes (Figure 1C). Changes could be observed for both high expressed genes as well as lower abundance transcripts with comparable numbers of up- and down-regulated transcripts in each comparison.

Gene expression levels correlated well with the enrichment of H3K4me3 at the corresponding TSS (Figure 1D), indicating that the ChIP-seq could serve as an indicator of midbrain transcriptional activity.

### *Pttg1* is the only transcription factor with altered midbrain expression between all three mouse strains

Gene expression changes linked to complex traits have been suggested to be explained by both small cumulative effects of *cis*-regulatory variants across numerous genes, and by *cis-* regulatory variants at “peripheral master regulators” such as TFs that can in *trans* influence a number of co-regulated genes directly linked to the trait (6). To better understand whether the observed gene expression changes in the mouse midbrain transcriptomes could be due to variants affecting upstream TFs, we further examined TFs included among the differentially expressed genes (DEGs). We first overlapped the DEGs from the pair-wise comparisons of the strains and identified 53 genes to be differentially expressed between all three strains (Figure 2A). Moreover, we identified a total of 1292 genes to be shared between at least two of the pair-wise comparisons of the strains (Supplementary Table S2). These genes are clustered in Figure 2B according to their gene expression profiles across the three strains with comparable numbers of genes showing particularly abundant or low expression levels in one or another strain. Next we used a manually curated list of 950 TFs (20), 841 of which could be detected in the midbrain, and identified 5 genes coding for TFs *(Pttg1, Npas1, Hes5, Scand1*, and *Zfp658*) to be differentially expressed in at least one of the mouse strains (Figure 2B). Interestingly, the number of differentially expressed TFs was much smaller than the 36 TFs that could be expected among the DEGs just by chance (hypergeometric test, p = 2.03*10^-11^). This lack of variation among TFs indicates a tight control of TF gene expression, which may need to be kept within a narrow range to allow for proper cellular function in the midbrain. Among the five TFs, only *Pttg1* showed a significant difference and higher than 2-fold change between all three strains (Figure 2B and Supplementary Table S2). In detail, C57BL/6J midbrain samples showed an average *Pttg1* expression of 14 RPKM (Reads Per Kilobase Million) and DBA/2J samples an average expression of 2.5 RPKM while in A/J midbrains *Pttg1* expression was never higher than 1 RPKM (Figure 2B and Supplementary Table S2). Differential midbrain expression levels of *Pttg1* between different mouse strains was confirmed by RT-qPCR, with A/J showing a particularly low expression level (Supplementary Figure S1A and S1B). Moreover, the H3K4me3 signal from ChIP-seq analysis was clearly reduced at the TSS of *Pttg1* gene in A/J compared to C57BL/6J, while no differences were observed at the TSS of neighbouring genes *Slu7* and *Clqtnf2* (Figure 2C). In addition, the signal in A/J appeared comparable or lower than in DBA/2J, despite the overall enrichment in DBA/2J samples being weaker than in the other two strains. These results suggest that reduced expression of *Pttg1* in the midbrain of A/J is due to decreased transcription at the locus.

**Figure 2.**
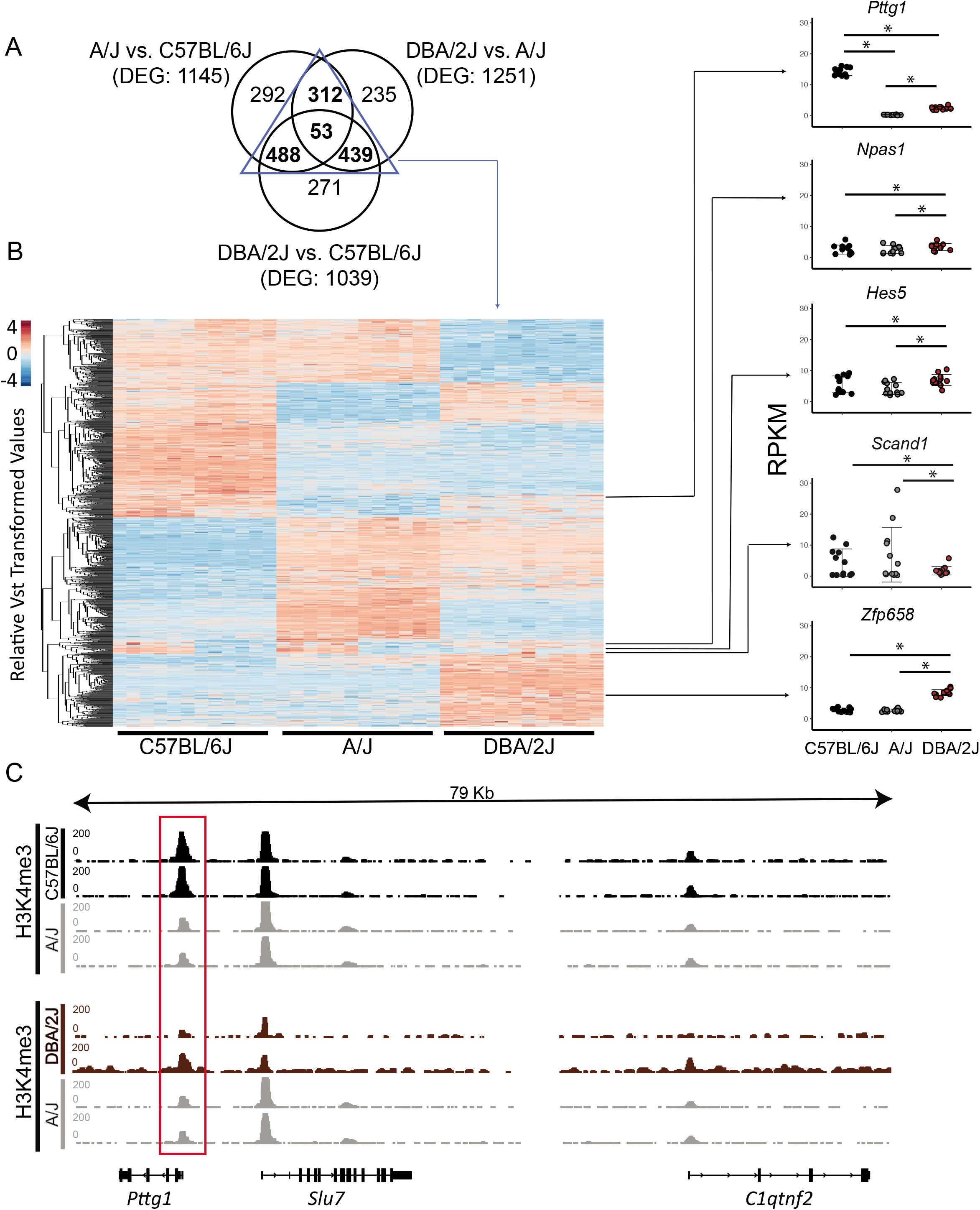
*Pttg1* is the only TF differentially expressed between the midbrains of 3-month-old C57BL/6J, A/J and DBA/2J mice. A. Venn diagram comparing DEGs of each pair-wise comparison of the mouse strains from Figure 1C. The majority of DEGs are shared by at least two comparisons. B. Heatmap of the expression of the 1292 DEGs shared between at least two of the comparisons. The read counts were vst-transformed and used for clustering. Expression levels of the five DEGs coding for TFs are shown as dot plots. *=FDR<0.05. C. The altered expression of *Pttg1* is accompanied by changes in H3K4me3 ChIP-seq signal at the *Pttg1* TSS. The H3K4me3 ChIP-seq was performed on two male replicates. The pair-wise comparisons (C57BL/6J vs. A/J and DBA2J vs. A/J) were performed by THOR with within-sample and between-sample normalizations. Normalized ChIP-seq signals are depicted in black (for C57BL/6J and DBA/2J) or in grey (for A/J). Red rectangle indicates *Pttg1* TSS.

Therefore *Pttg1* appears to be a prime candidate for explaining midbrain transcriptomic differences between the mouse strains.

### Loss of *Pttg1* leads to changes in the midbrain transcriptome during aging

Given that *Pttg1* encodes the only TF with significantly altered expression levels between all three mouse strains, we investigated the role of midbrain PTTG1 in more detail. To test whether altered expression of *Pttg1* alone can indeed influence the midbrain transcriptome, we investigated C57BL/6J congenic *Pttg1^-/-^* mice. In contrast to differences in A/J or DBA/2J, deletion of *Pttg1* in the 3-month old mice leads to minor transcriptomic changes with 3 additional genes differentially expressed compared to the *Pttg1*^+/+^ littermates (Figure 3A). Two of these *(Thg11* and *Ubilcp1*) were previously found to strongly correlate with *Pttg1* expression across different mouse strains, and to be genetically associated with neocortex volume (21), while the third gene *(Gm12663)* is an anti-sense transcript of *Ubilcp1.* Moreover, the expression of these genes is dependant of *Pttg1* expression level when corroborating the analysis with 3-month-old *Pttg1^+/-^*, with *Ubilcp1* showing positive, and *Thg11* and *Gm12663* showing negative correlation with *Pttg1* levels (Figure 3B).

**Figure 3.**
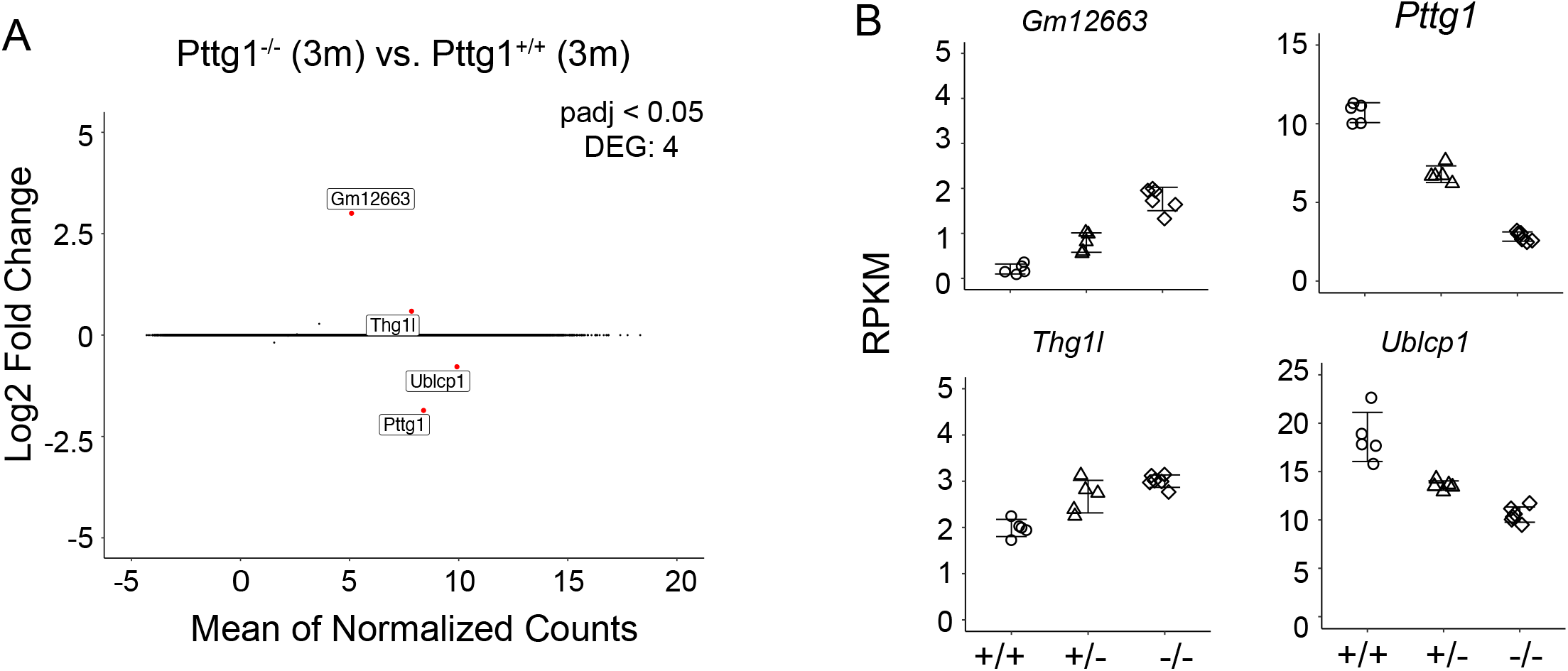
Loss of *Pttg1* leads to minimal changes in the midbrain transcriptome in 3-month old mice. A. RNA-seq analysis identifies four DEGs in comparison of the congenic C57BL/6J *Pttg1_-/-_* vs. *Pttg1^+/+^* mice at the age of 3 months. MA plot was generated as in Figure 1C with labelling of the four DEGs *(Pttg1, Thg11, Ublcp1, Gm12663)* that are indicated as red dots. B. The expression of *Ublcp1* is positively correlated with *Pttg1* across genotypes, while *Gm12663* and *Thg11* show negative correlation with *Pttg1.* The dot plots indicate the expression levels of the DEGs as RPKM in isolated midbrains of *Pttg1^+/+^*, *Pttg1^+/-^*, and *Pttg1^-/-^* mice.

Although the observed midbrain transcriptome changes in *Pttg1^-/-^* mice were minimal, we were curious to elucidate whether these early changes would lead to additional transcriptomic differences at an older age. We therefore performed further RNA-seq analysis with isolated midbrains of a cohort of six aged mice from each C57BL/6J, A/J, and DBA/2J strains (all 9 months old), and C57BL/6J congenic *Pttg1^-/-^* mice (9-13 months old) (Figure 4A). Interestingly, comparing samples from 9-month-old wild-type C57BL/6J or A/J to those from younger 3-month-old mice of the respective strains identified almost no genes with strong expression changes of 5-fold or more (log_2_FC>2.25) (Figure 4B and Supplementary Table S3). Similarly, comparison of 9-month-old DBA/2J midbrain transcriptome to the younger counterparts revealed only 57 strongly altered genes. Conversely, the midbrain samples of *Pttg1^-/-^* mice showed over 300 genes that were strongly differentially expressed in the aged mice compared to 3-month-old mice, as shown in the Vulcano plot in Figure 4B.

**Figure 4.**
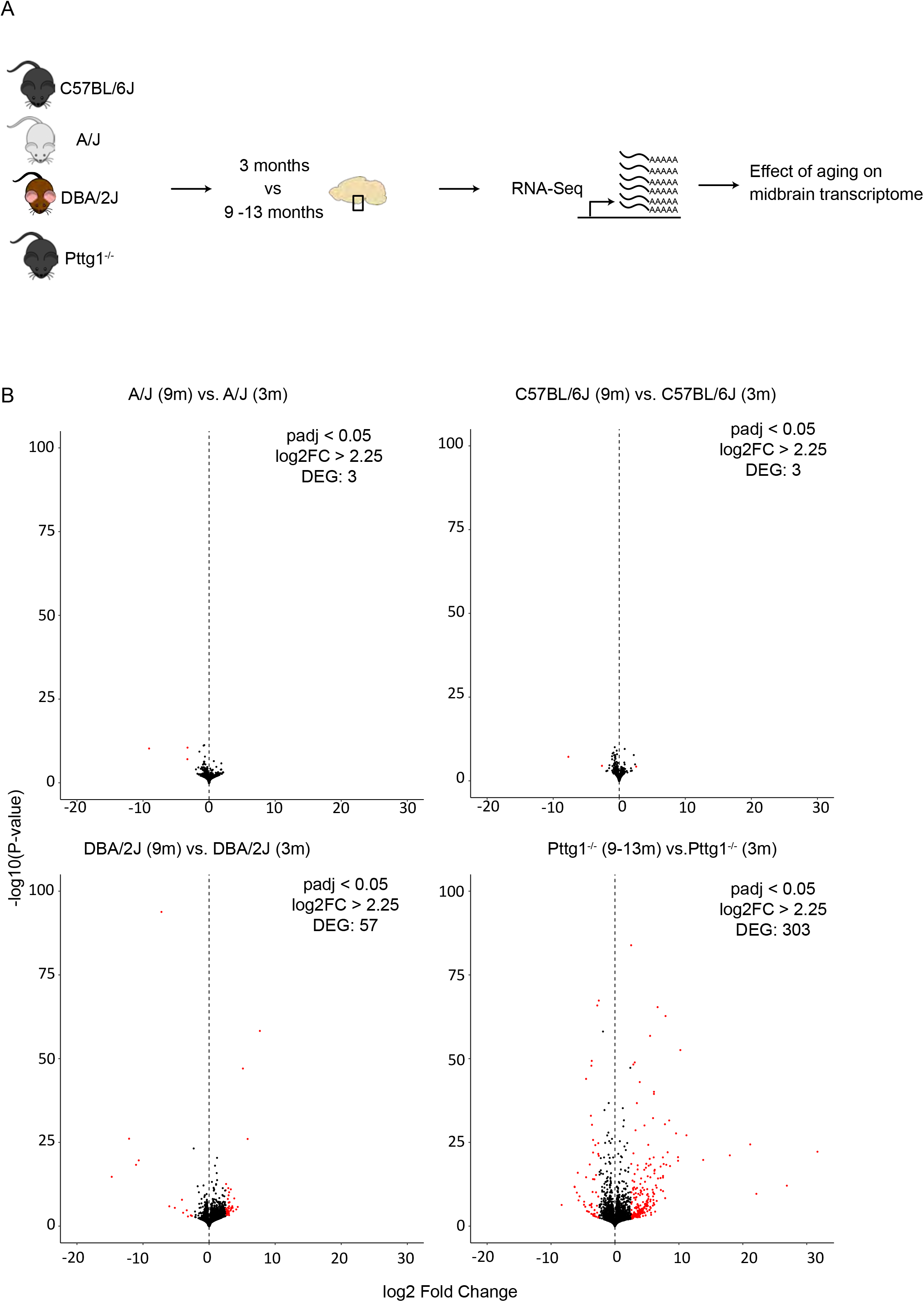
Loss of *Pttg1* leads to significant transcriptomic changes in the midbrain during aging. A. Schematic representation of the experimental set-up. The ventral midbrains of 9-month-old C57BL/6J, A/J and DBA/2J mice, and 9-13-month-old congenic C57BL/6J *Pttg1^-/-^* mice were used for RNA-seq as in Figures 1 and 3. B. Comparison of midbrain transcriptome of 9-month-old mice to the midbrain transcriptome of the corresponding strains at 3 months of age. *Pttg1* deletion leads to more significant and higher gene expression changes than observed for wild-type mouse strains during aging. Vulcano plots from left to right: A/J, C57BL/6J, DBA/2J, and congenic C57BL/6J Pttg1^-/-^. The x-axis represents the mean log_2_-fold change for all replicates and the y-axis represents the significance of change as −log_1_0 (p-value). Each dot represents one gene. Genes with FDR<0.05 and log_2_-fold change (log_2_FC)>2.25 are indicated in red and referred to as DEGs.

### *Pttg1* contributes to gene regulatory variation in the midbrain cell types during aging

To obtain a broader overview of the extent and the direction of transcriptomic changes across the studied mouse strains and ages, we performed PCA analysis for all 78 midbrain transcriptome profiles. Interestingly, the PCA revealed that over half of the variance between the studied mice was explained by the first and the second principle components (PCs) that separated the mice according to genetic background (Figure 5A). C57BL/6J mice were separated from A/J and DBA/2J along PC1 while A/J and DBA/2J were separated from each other along PC2. Consistent with the small number of DEGs in 3-month-old *Pttg1^-/-^* mice, they clustered closely together with their heterozygous *Pttg1^+/-^* littermates, and with wild-type C57BL/6J mice. Also, aged mice clustered largely together with their genetically identical counterparts for each A/J, DBA/2J, and C57BL/6J. However, for the aged cohort of *Pttg1^-/-^* mice, the transcriptome profiles had significantly shifted along PC1 from C57BL/6J towards A/J and DBA/2J (Figure 5A).

**Figure 5.**
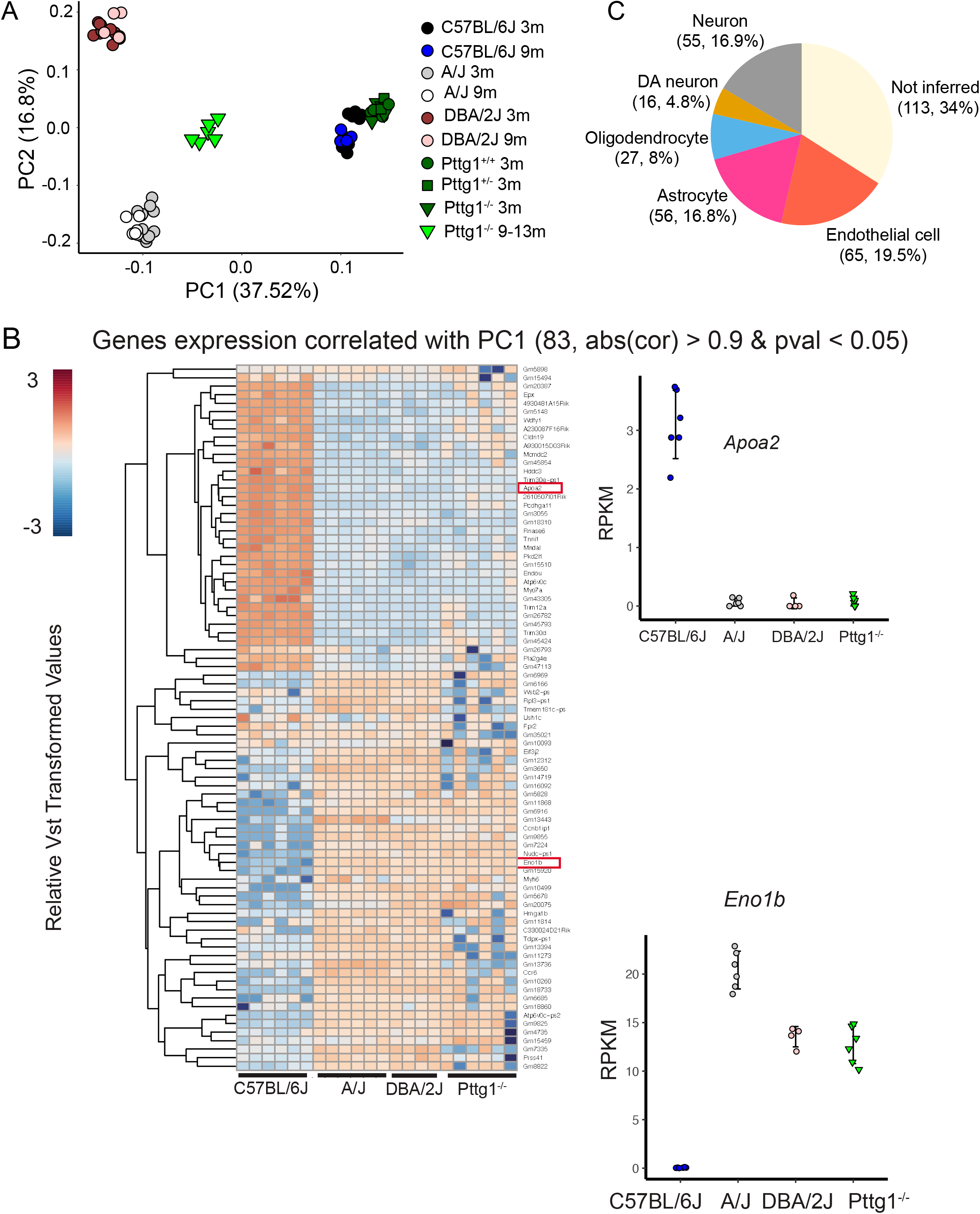
C57BL/6J *Pttg1^-/-^* midbrain transcriptome shift towards A/J during aging. A. Principle component analysis showing transcriptome level differences in the midbrains of C57BL/6J, DNA/2J, and A/J mice at the age of 3 and 9 months, congenic C57BL/6J *Pttg1^+/+^*, *Pttg1^+/-^*, and *Pttg1^-/-^* at 3 months, and *Pttg1^-/-^* at 9 to 13 months. Individual mice are indicated with black circles (C57BL/6J 3m), blue circles (C57BL/6J 9m), grey circles (A/J 3m), white circles (A/J 9m), brown circles (DBA/2J 3m), light brown circles (DBA/2J 9m), green circles *(Pttg1^+/+^* 3m), dark green rectangles *(Pttg1^+/-^* 3m), dark green triangles (*Pttg1^-/-^* 3m), or light green triangles *(Pttg1^-/-^* 9-13m). No gender bias was observed. B. Heat map of the differential genes associated with principle component 1 in panel A. Gene expression profile of *Pttg1^-/-^* mice clusters with A/J and DBA/2J mice instead of C57BL/6J. C. Deconvolution of differential gene expression using single cell RNA-seq was done for 331 genes contributing the most to PC1 in panel A and detected as expressed in 5 major cell types of the scRNA-seq data. The number of genes and their proportion of all analysed genes are shown for each cell type.

Analysis for genes that contributed most to the differences along PC1 revealed genes that were altered not only in A/J and DBA/2J strains but also in aged *Pttg1^-/-^* mice when compared to C57BL/6J (Figure 5B). Furthermore, gene changes in *Pttg1^-/-^*, A/J, and DBA/2J mice showed the same directionality, with the *Pttg1^-/-^* mice clustering together with A/J and DBA/2J rather than C57BL/6J when analysed with hierarchical clustering.

Finally, to see whether the loss of *Pttg1* was specifically affecting only some of the cell types in the midbrain, we performed deconvolution analysis of the DEGs contributing to PC1 using mouse midbrain single cell RNA-seq data (22). Based on the inference, the DEGs included genes preferentially expressed in many different cell types, including different types of neurons such Th^+^ DAns, oligodendrocytes, astrocytes, and endothelial cells (Figure 5C). Additionally, over a third of the genes could not be inferred, suggesting they are expressed broadly across multiple different cell types.

Taken together, the results indicate that loss of *Pttg1* leads to only limited transcriptomic changes in the midbrain of young mice, but can lead to substantial differences during aging, with parts of C57BL/6J transcriptome shifted towards A/J and DBA/2J in aging mice. Thus, our data indicate that PTTG1 contributes to transcriptome differences in multiple cell types of the midbrain between the three genetically diverse mouse strains.

## Discussion

We investigated gene expression differences between mouse strains to understand how genetic variation can influence midbrain and its important cell types such as DAns that control motor function and behaviour. Our transcriptomic analysis revealed extensive changes in midbrain gene expression between the 3 mouse strains and highlighted *Pttg1* as an important regulator of midbrain transcriptome during aging.

The observed midbrain transcriptomic differences are comparable to the transcriptome level changes observed between mouse strains in other tissues such as lung (23), striatum (24), and retina (25), or in specific cell types such as macrophages (26) and other immune cells (27). Interestingly, despite the obvious variation in the gene expression between the mouse strains, genes encoding for TFs are under-represented among the DEGs in the mouse midbrain. This finding is consistent with similar results from plants (28), where TF coding genes were also found to be under-represented among the genes showing differential expression between genetically diverse strains. Such findings are likely to be due to natural selection against phenotypes arising from major variation in TF expression levels that could be detrimental for the normal functioning of an organism.

The only TF showing significant changes in the ventral midbrain between all three mouse strains is *Pttg1*, also known as securin. *Pttg1* was originally described as an oncogene in pituitary tumors (29) and found to regulate sister chromatid adhesion in M-phase of cell cycle (30). However, the protein has multiple functions and also a role as a DNA-binding transcriptional activator has been described (reviewed in Vlotides *et al.*, 2007 (31)).

Little is known about the neurological functions of PTTG1. Keeley *et al.*, 2014 (32) identified a link between PTTG1 and the central nervous system, showing increased *Pttg1* expression in retinas of C57BL/6J mice compared to A/J due to a *cis* deletion variant, consistent with our findings in the midbrain. Interestingly, differential *Pttg1* expression correlated with mosaic regularity variation across 25 recombinant inbred strains derived from the two parental C57BL/6J and A/J mouse strains, involving PTTG1 in the patterning of a type of retinal neurons, the amacrine cells. Moreover, *Pttg1* expression in neocortex correlates with neocortical volume and the locus is genetically associated with this trait (21). Therefore, *Pttg1* appears to play a role in development or maintenance of central nervous system, and our results indicate its possible involvement in genetic control of midbrain cell types. Indeed, previous work using microarrays found >1400 genes to be misregulated across the whole brain of *Pttg1^-/-^* mice at the age of 3-5 months (33). While we identified far fewer DEGs specifically in the midbrain of the 3 month-old *Pttg1^-/-^* mice at our significance cut-off (FDR<0.05) using RNA-seq, this increased increased significantly during aging. Importantly, the overall transcriptomic profile shifted towards the profiles of A/J and DBA/2J (Figure 5), indicating that *Pttg1* might indeed exhibit genetic control over gene expression in the midbrain, although additional genetic factors are likely altered to contribute to these changes already in young mice.

It has been previously reported that *Pttg1* is involved in many biological functions such as regulation of sister chromatid separation, DNA repair or senescence processes (30, 34, 35). Interestingly, a deconvolution analysis of the gene expression changes using single cell RNA-seq analysis indicated that the loss of *Pttg1* influenced gene expression across multiple cell types. However, these changes become observable only during aging. Unlike human brain, mouse brain volume has been shown to increase still during adulthood between 6 and 14 months of age (36). Given the abovementioned role of *Pttg1* in regulation of neocortex volume and its effect of gene expression in multiple cell types during aging, it is tempting to speculate that *Pttg1* would contribute also to control of midbrain volume. Interestingly, a greater brain volume has been reported for C57BL/6J than A/J (37, 38).

## Conclusions

Rather than being entirely explained by the TF expression levels due to *cis*-variation at the *Pttg1* locus, complex traits like midbrain gene expression could be due to cumulative *cis*- and *trans*-regulatory variants across TF binding sites controlling the DEGs. In the future, mapping QTLs associated with the DAn’s traits across mouse, together with the transcriptomic and epigenomic data generated as part of this work, will enable the identification of further regulatory variants and their impact on midbrain expression phenotype and function of the nigrostriatal circuitry. While linking complex traits such as behaviour and motor function to specific gene expression changes will require further studies, our work highlights the role of *Pttg1* as regulator of mouse midbrain gene expression phenotype and paves way for further identification of additional genetic regulators.

## Methods

### Animals

All experiments were performed in accordance with the European Communities Council Directive 2010/63/EU, approved by appropriate government agencies and respecting the 3 Rs’ requirements for Animal Welfare. For the mice bred in the Animal Facility of University of Luxembourg, all experiments in mice were performed according to the national guidelines of the animal welfare law in Luxembourg (*Règlement grand-ducal* adopted on January 11^th^, 2013). The protocol was reviewed and approved by the Animal Experimentation Ethics Committee (AEEC). For the mice bred in Helmholtz Centre for Infection Research (Braunschweig, Germany), all experiments were performed according to the national guidelines of the animal welfare law in Germany (BGBl. I S. 1206, 1313 and BGBl. I S. 1934). The protocol was reviewed and approved by the ‘Niedersächsisches Landesamt für Verbraucherschutz und Lebensmittelsicherheit, Oldenburg, Germany’ (Permit Number: 33.9-42502-05-11A193). Mice were housed on a 12 hours-light/dark cycle and provided food and water *ad libitum.* Three mouse strains, C57BL6/6J, A/J and DBA/2J, were used in this study. C57BL/6J and DBA/2J mice were purchased from the provider of Jackson Laboratory in Europe (Charles River). Study cohorts were either directly used after a 2-weeks resting period or were bred in house. The A/J breeders were directly purchased from Jackson Laboratory and the study cohorts were bred either at the Helmholtz Centre for Infection Research (Braunschweig, Germany) or in-house at the Animal Facility of University of Luxembourg (Esch-sur-Alzette, Luxembourg). The *Pttg1* knock-out transgenic line was established at Cedars Sinai Medical Center (39) and *Pttg1^-/-^* mice had been backcrossed to C57BL/6J for more than 10 generations. 9-to 13-month old *Pttg1^-/-^* female mice were bred at the Cedars Sinai Medical Center (Los Angeles, USA). The *Pttg1^+/-^* mice were bred in-house to generate a 3 month-old study cohort (*Pttg1^+/+^, Pttg1^+/-^*, and *Pttg1^-/-^*) and to maintain a colony at the local animal facility. Brain samples from the 9 to 13 month-old *Pttg1^-/-^* mice were collected as described below and 9 month-old strain-matched C57BL/6J were used as a control.

At each age group, 5 to 17 mice were anesthetized with a ketamine-medetomidine mix (150 and 1 mg/kg, respectively) and intracardially perfused with PBS (phosphate-buffered saline) before extracting the brain. One hemibrain of each mouse was dissected for midbrain which was immediately snap-frozen, stored at −80°C, and used for qPCR, RNA-seq, and ChIP-seq analysis as described below.

### RT-qPCR

The RNA expression of genes of interest was measured in the midbrains of C57BL/6J, A/J, and DBA/2J. RNA was extracted from the midbrain of each mouse using the RNeasy^®^ Plus Universal Mini Kit (Qiagen, Germany). The reverse transcription was performed using 300 ng of total RNA mixed with 3.8 μM of oligo(dT)20 (Life Technologies) and 0.8 mM of dNTP Mix (Invitrogen). After heating the mixture to 65°C for 5 minutes and an incubation on ice for 1 min, a mix of first-strand buffer, 5 mM of DTT (Invitrogen), RNAse OUT™ (Invitrogen) and 200 units of SuperScript III reverse transcriptase (200 units/μL, Invitrogen) was added to the RNA. The mixture was incubated at 50°C for 60 minutes and then the reaction was inactivated by heating at 70°C for 15 minutes. After adding 80 μL of RNAse free water, the cDNA is stored at −20°C.

RT-qPCR was performed to measure the RNA expression of several genes using the Applied Biosystems 7500 Fast Real-Time PCR System. Each reaction had 5 μL of cDNA, 5 μL of primer mixture (forward and backward primers) (2 μM) and 10 μL of the Absolute Blue qPCR SYBR Green Low ROX Mix (ThermoFisher Scientific, AB4322B). The conditions of the PCR reaction were the following: 95°C for 15 minutes and repeating 40 cycles of 95°C for 15 seconds, 55°C for 15 seconds and 72°C for 30 seconds. The gene expression level was calculated using the 2^-(ΔΔCt)^ method. The ΔΔCt refers to ΔCt_(target gene)_ – ΔCt_(housekeeping gene)test_ – (ΔCt_(target gene)_ – ΔCt_(housekeeping gene)_)_control_. The sequences of the used primers are provided in the Supplementary Table S1.

### RNA-seq

The RNA sequencing of 6 C57BL/6J and 6 A/J samples from both 3 month-old and 9-month old mice was done at the sequencing platform of the Genomics Core Facility in EMBL Heidelberg, Germany. The samples were processed by Illumina CBot. The single-end, stranded sequencing was applied by the Illumina NextSeq 500 machine with read length of 80 bp.

The remaining RNA-seq samples were processed at the sequencing platform in the Luxembourg Centre for Systems Biomedicine (LCSB) of the University of Luxembourg. The RNA quality was determined by Agilent 2100 Bioanalyzer and the concentration was quantified by Nanodrop. The TruSeq Stranded mRNA Library Prep kit (Illumina) was used for library preparation with 1 μg of RNA as input according to the manufacturer’s instructions. The libraries were then adjusted to 4 nM. The single-end, stranded sequencing was applied by the Illumina NextSeq 500 machine with read length of 75 bp.

The raw reads quality was assessed by FastQC (v0.11.5) (40). Using the PALEOMIX pipeline (v1.2.12) (41), AdapterRemoval (v2.1.7) (42) was used to remove adapters, with a minimum length of the remaining reads set to 25 bp. The rRNA reads were removed using SortMeRNA (v2.1) (43). After removal of adapters and rRNA reads, the quality of the files was re-assessed by FastQC. The mapping was done by STAR (v.2.5.2b) (44). The mouse reference genome, GRCm38.p5 (mm10, patch 5), was downloaded from GENCODE. The suit tool Picard (v2.10.9) (45) validated the BAM files. Raw FASTQ files were deposited in ArrayExpress with the accession number E-MTAB-8333.

The reads were counted using *featureCounts* from the R package *Rsubread* (v1.28.1) (46). The DEGs were called using R package *DESeq2* (v1.20.0) (47). RPKM for each gene in each sample was calculated as reads divided by the scale factor and the gene length (kb). The scale factor was calculated as library size divided by 1 million.

### Chromatin Immunoprecipitation (ChIP)

ChIP was performed on the dissected mouse midbrain tissue. The fresh tissue was snap frozen for at least a week before crosslinking with formaldehyde (Sigma-Aldrich, F8775-25ML) at a final concentration of 1.5% in PBS (Lonza, BE17-516F) for 10 minutes at room temperature. The formaldehyde was quenched by glycine (Carl Roth, 3908.3) at a final concentration of 125 mM for 5 minutes at room temperature, followed by centrifugation at 1,300 rpm for 5 minutes at 7°C. The fixed tissue was washed twice for 2 minutes with icecold PBS plus 1x cOmplete™ mini Proteinase Inhibitor (PI) Cocktail (Roche, 11846145001). The tissue was minced by the Dounce Tissue Grinder (Sigma, D8939-1SET), the lysate of which was centrifuged at 1,300 rpm for 5 minutes at 7°C. The pellet was suspended in icecold Lysis Buffer [5 mM 1,4-piperazinediethanesulfonic acid (PIPES) pH8.0 (Carl Roth, 9156.3), 85 mM potassium chloride (KCl) (PanReac AppliChem, A2939), 0.5% 4-Nonylphenyl-polyethylene glycol (NP-40) (Fluka Biochemika, 74385)] with 1xPI, and kept on ice for 30 minutes. The tissue lysate was centrifuged 2,500 rpm for 10 minutes at 7°C. The pellet was suspended with ice-cold Shearing Buffer [50mM Tris Base pH 8.1, 10 mM ethylenediamine tetraacetic acid (EDTA) (Carl Roth, CN06.3), 0.1% sodium odecylsulfate (SDS) (PanReac Applichem, A7249), 0.5% sodium deoxycholate (Fluka Biochemika, 30970)] with 1x PI.

The sonication (Diagenode Bioruptor^®^ Pico Sonication System with minichiller 3000) was used to shear the chromatin with program 30 seconds on, 30 seconds off with 35 cycles at 4°C. After sonication the cell debris was removed by centrifugation at 14,000 rpm for 10 minutes at 7°C. The concentration of the sheared and reverse crosslinked chromatin was measured by Nanodrop 2000c (Thermo Scientific, E597) and shearing was confirmed to produce chromatin fragments of 100 bp to 200 bp.

Each reaction had 10 – 14 μg of chromatin, of which 10% of the aliquot was used as input DNA. The chromatin sample was diluted 1:10 with Modified RIPA buffer [140 mM NaCl (Carl Roth, 3957.2), 10 mM Tris pH 7.5, 1 mM EDTA, 0.5 mM ethylene glycol-bis-N,N,N’,N’-tetraacetic acid (EGTA) (Carl Roth, 3054.3), 1% Triton X-100, 0.01% SDS, 0.1% sodium deoxycholate] with 1x PI, followed by addition of 5 μL of H3K4me3 (histone H3 lysine 4 trimethylation) antibody (Millipore, 17-614) and incubation overnight at 4°C with rotation. After incubation, the immunocomplexes were collected with 25 μL of PureProteome™ Protein A Magnetic (PAM) Beads (Millipore, LSKMAGA10) for 2 hours at 4°C with rotation.

The beads were washed twice with 800 μL of Wash Buffer 1 (WB1) [20 mM Tris pH 8.1, 50 mM NaCl, 2mM EDTA, 1% Triton X-100, 0.1% SDS], once with 800 μL of Wash Buffer 2 (WB2) [10 mM Tris pH 8.1, 150 mM NaCl, 1 mM EDTA, 1% NP-40, 1% sodium deoxycholate, 250 mM lithium chloride (LiCl) (Carl Roth, 3739.1)], and twice with 800 μL of Tris-EDTA (TE) Buffer [10 mM Tris PH 8.1, 1 mM EDTA pH 8.0]. The beads were resuspended in 100 μL of ChIP Elution Buffer [0.1 M sodium bicarbonate (NaHCO_3_) (Sigma-Aldrich, S5761)and 1 % SDS]. After the elution, the chromatin and the 10% input were both reverse-crosslinked at 65°C for 3 hours with 10 μg of RNase A (ThermoFisher, EN0531) and 20 μg of thermoresistant proteinase K (ThermoFisher, EO0491), followed by purification with MiniElute Reaction Cleanup Kit (Qiagen, 28206) according to the manufacture’s instruction.

The concentration of the chromatin was measured by Qubit^®^ dsDNA HS Assay Kit (ThermoFisher, Q32851) and Qubit 1.0 fluorometer (Invitrogen, Q32857) according to the manufacturer’s instructions and rest of the chromatin was used for high-throughput sequencing.

### ChIP-seq

The sequencing of the chromatin samples was done at the sequencing platform in the LCSB of the University of Luxembourg. The single-end, unstranded sequencing was applied by the Illumina NextSeq 500 machine with read length of 75 bp. The raw reads quality was assessed by FastQC (v0.11.5) (40). The PALEOMIX pipeline (v1.2.12) (41) was used to generate BAM files from the FASTQ files, including steps of adapter removal, mapping and duplicate marking. The mapping was done by BWA (v.0.7.16a) (48), with backtrack algorithm using the quality offset of Phred score to 33. Duplicate reads were marked but not discarded. The mouse reference genome, GRCm38.p5 (mm10, patch 5), was downloaded from GENCODE (https://www.gencodegenes.org/). The suit tool Picard (v2.10.9) (45) was used to validate the BAM files. Raw FASTQ files were deposited in ArrayExpress with the accession number E-MTAB-8333.

The H3K4me3 ChIP-seq peaks were called by Model-based analysis of ChIP-seq (MACS, v2.1.1) (49). The signal normalization in pairwise comparison was done by THOR (v0.10.2) (50), with TMM normalization and adjusted p-value cut-off 0.01.

### Principle Component Analysis (PCA)

The raw counts were normalized to library size and log_2_-transformed using *DESeq2* (v1.20.0). The PCs were calculated with 500 genes which have the most varied expression across samples.

### Bulk RNA-seq data deconvolution using single cell RNA-seq data

The bulk RNA-seq deconvolution was done with CIBERSORTx (https://cibersortx.stanford.edu/) (51). The signature matrix on SN of single cell RNA-seq was constructed with DropViz (http://dropviz.org/) with default parameters. The expression of 332 genes correlating with PC1 (pvalue < 0.05) from Figure 5A in 5 cell types (neuron, dopaminergic neuron, oligodendrocyte, astrocyte, endothelial cells) were inferred with default parameters.

### Statistical Analysis

The p-value of DEGs called from pair-wise comparisons in RNA-seq was adjusted for multiple testing with the Benjamini-Hochberg procedure with cutoff below 0.05. The significance for peak calling by MACS2 in ChIP-seq experiments was detailed in Zhang et al., 2008 (27). The significance in ChIP-seq signal normalization was defined with multiple test correction (Benjamini/Hochberg) for p-values with cutoff below 0.05.

## Abbreviations

DAns: dopaminergic neurons
DEGs: differentially expressed genes
EDTA: ethylenediamine tetraacetic acid
H3K4me3: histone H3 lysine 4 trimethylation
LCSB: Luxembourg Centre for Systems Biomedicine
log_2_FC: log_2_-fold change
NP-40: 4-Nonylphenyl-polyethylene glycol
PC: principle component
PCA: principle component analysis
PD: Parkinson’s disease
PI: Proteinase Inhibitor
*Pttg1*: Pituitary Tumor Transforming Gene 1
RPKM: Reads Per Kilobase Million
SDS: sodium odecylsulfate
SN: substantia nigra
TFs: transcription factors
TSS: transcription start sites
VTA: ventral tegmental area.

## Acknowledgements

We would like to thank Drs Aurélien Ginolhac and Anthoula Gaigneaux for their support with bioinformatic analysis and EMBL Gene Core at Heidelberg for support with high-throughput sequencing, and Dr Djalil Coowar (Animal Facility of University of Luxembourg) for help with breeding of experimental mice. KS would like to thank the animal caretakers at the Central Animal Facilities of the HZI for maintaining the mice. The computational analysis presented in this paper were carried out using the HPC facilities of the University of Luxembourg (51).

