## Supplementary figures for "*Pituitary Tumor Transforming Gene 1* orchestrates gene regulatory variation in mouse ventral midbrain during aging"

#### Supplementary Information

### Supplementary Figures

**Supplementary Figure S1. RT-PCR measurements of *Pttg1* expression in isolated midbrains are consistent with the RNA-seq results.**

A. *Pttg1* expression measured by RT-PCR is consistent with the RNA-seq data across the three strains. Expression levels are presented relative to *Gapdh*. Two-sided Student's t-test was used for statistical testing. *=p<0.05.

B. *Pttg1* expression measured by RT-PCR is consistent with the RNA-seq data across the 3 months old $Pttg1^{+/+}$, $Pttg1^{+/-}$, $Pttg1^{-/-}$, and 9-13 months old $Pttg1^{-/-}$ mice. Expression levels are presented relative to *Rpl13a*. Two-sided Student's t-test was used for statistical testing. *=p<0.05.

Supplementary Figure S1

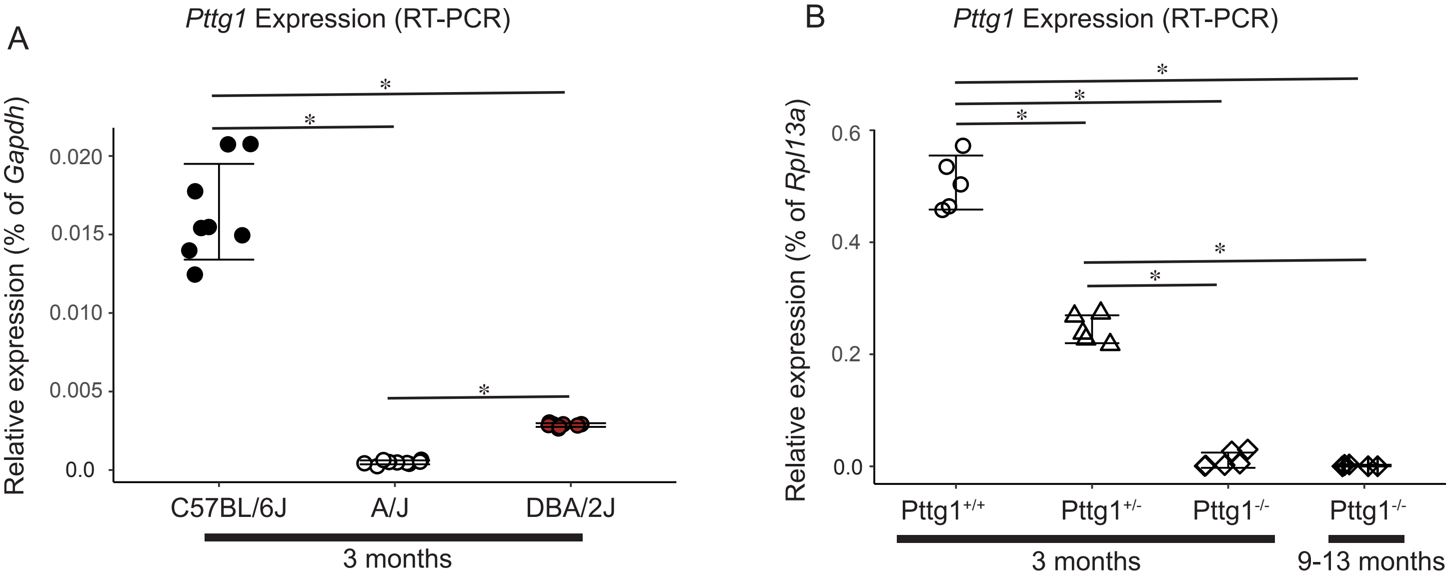

### Supplementary Tables

**Supplementary Table S1**. **Primer sequences used in the study.**

| **Gene** | **Forward primer (5‘ – 3‘)** | **Reverse primer (5‘ – 3‘)** |
|---|---|---|
| *Pttg1* | TCAAGGTCGGCTGTTTTGGT | AGTTGCCGAAAAGCCTATGAAG |
| *Rpl13a* | TGGTCCCTGCTGCTCTCA | CCCCAGGTAAGCAAACTTTCT |
| *Gapdh* | TGCGACTTCAACAGCAACTC | CTGCTCAGTGTCCTTGCTG |

**Supplementary Table S2. DEGs (FDR < 0.05, log2FC > 1) shared by at least two comparisons in Figure 2B.** The base mean, log2FC, and FDR are reported for each gene in each comparison: A/J vs. C57BL/6J: 853 genes; DBA/2J vs. A/J: 804 genes; DBA/2J vs. C57BL6/J: 980 genes.

**Supplementary Table S3. DEGs (FDR < 0.05, log2FC > 2.5) from 3 months old vs. 9 months old mice.** The DEGs from C57BL/6J A/J, DBA/2J and *Pttg1*$^{-/-}$ mice are shown on individual worksheets. The base mean, log2FC, and FDR are reported for each gene in each comparison.
